# Aging alters tumor cell - T cell crosstalk to promote breast cancer progression

**DOI:** 10.1101/2025.11.17.688754

**Authors:** Shanshan Yin, Kay T Yeung, Sainath Mamde, Kate Lin, Armin Gandhi, Xue Lei, Andrew Davis, Rouven Arnold, Jing Yang, Peter D. Adams

## Abstract

Age is a dominant risk factor for all major breast cancer subtypes. However, the mechanisms by which aging influences tumor development remain unclear. Using a novel mouse model whereby breast cancer is induced *in situ* in young and old wild-type mice via intraductal delivery of a lentivirus encoding the *HER2/neu* oncogene, we found that old mice exhibited a higher oncogene-induced tumor burden than young mice. Old tumor cells showed reduced expression of interferon-related genes, particularly the T cell-recruiting chemokines *Cxcl9* and *Cxcl10*, linked to their altered chromatin accessibility. *CXCL9/10* expression also declined with age in human HER2+ tumors. Correspondingly, old tumors exhibited fewer T cells within tumor lesions. Targeted interventions showed that decreased expression of *Cxcl9*/*10* is responsible for reduced T cell infiltration and weakened anti-tumor immunity. These results show how aged tumor cells are impaired in their recruitment of immune cells, leading to a defective anti-tumor immune response.

## Introduction

Aging is the biggest risk factor for cancer. Progressive accumulation of genetic mutations over time contributes to the age-associated increase in cancer incidence^1^. However, although cells do acquire mutations with age, cancer in the elderly is not likely explained solely by the accumulation of somatic mutations^2,3^. Several of the hallmarks of aging, such as epigenetic alterations^4^, immune dysregulation^5^, metabolic remodeling^6^, and cellular senescence^7–10^, are shared between aging and cancer and likely collectively contribute to increased cancer susceptibility with age^11,12^.

Breast cancer is the most frequently diagnosed malignancy among women worldwide. Like most adult human cancers, age is a major risk factor for breast cancer. According to the NCI, women at age 60 have a 7-fold higher relative breast cancer risk than women at age 30, and more than 50% of patients are over 60 years of age at diagnosis^13,14^. The incidence of all three main subtypes of breast cancer, ER/PR+, HER2+ and triple negative, increases with age^14,15^. Older breast cancer patients also have higher mortalities and poorer survival outcomes^16–18^.

Mammary tissues are comprised of epithelial cells and the surrounding microenvironment of immune cells and stromal cells^19^. The luminal epithelial cells are considered to be the cell of origin of breast cancer^20^. Mammary epithelial cells undergo age-associated shifts in cell composition and gene expression linked to cancer susceptibility^21–23^. Aging also weakens epithelial identity and function^23^, expands stem-cell-like populations with increased transformation potential^24,25^, and promotes a late senescence phase enriched for cancer-associated pathways^26^. The mammary microenvironment similarly exhibits age-related, cancer promoting changes^27^. In rat, naïve B and T cells in the mammary tissues decline in number^25^. In mice, aging attenuates interferon signaling and reduces the efficacy of immune checkpoint blockade against triple-negative breast cancer^28^, promotes metastasis through fibrotic remodelling^29^, and induces invasive cellular phenotypes in vitro^30^. Together, both epithelial-intrinsic and microenvironmental factors contribute to the age-dependent increase in mammary cancer susceptibility. However, how aging affects epithelial–microenvironmental crosstalk and its role in tumor development remains to be fully elucidated.

Contributing to our lack of knowledge of the role of aging in cancer, there are few mouse models that fully recapitulate the age-dependence of breast cancer or other cancers. Studies employing tumor cell transplants into young and old mice have been very informative on the role of the young and old tissue microenvironment affecting primary tumors and metastasis^28,31–33^. However, transplant models do not inform on the role of aging in tumor initiating epithelial cells or tumor cells, since the transplanted tumor cells are identical.

Here, we sought to address this issue with a complementary mouse model. In this model, the *HER2/neu* oncogene is introduced directly into young and old mammary epithelial cells within their native tissue environment^34–36^. Compared to tumor cell transplant models, this approach enables the study of tumorigenesis in an intact microenvironment, providing a powerful platform to investigate how aging influences tumor cells and their interactions with microenvironmental components. This model thus offers new insights into the age dependence of breast tumorigenesis.

## Results

### Old mice are prone to a higher breast tumor burden

Based on aforementioned data showing substantial cancer-relevant changes with age in mammary tissues, we hypothesized that young and old mammary glands would respond differently, in terms of tumor initiation and/or tumor progression, to the same activated oncogene. To test this hypothesis, we adopted a method whereby a lentivirus encoding an activated truncated rat *Erbb2* (*HER2/neu*) oncogene is intraductally injected into mammary ducts of female BALB/c mice^37^. This method has been used in young mice^34–36^ to model HER2+ breast cancer, of which the incidence is >5x higher in women ages 65-74 than in women around 30 years of age^14^ (Figure 1A). To adapt this model to understand the effect of age, we used mice 3-5 months of age as “young” mice, equivalent to ∼25 years in humans, and 18-20-months as “old” mice, equivalent to ∼60 years in humans. The same amount of virus was injected into the mammary ducts of young and old mice.

**Figure 1:**
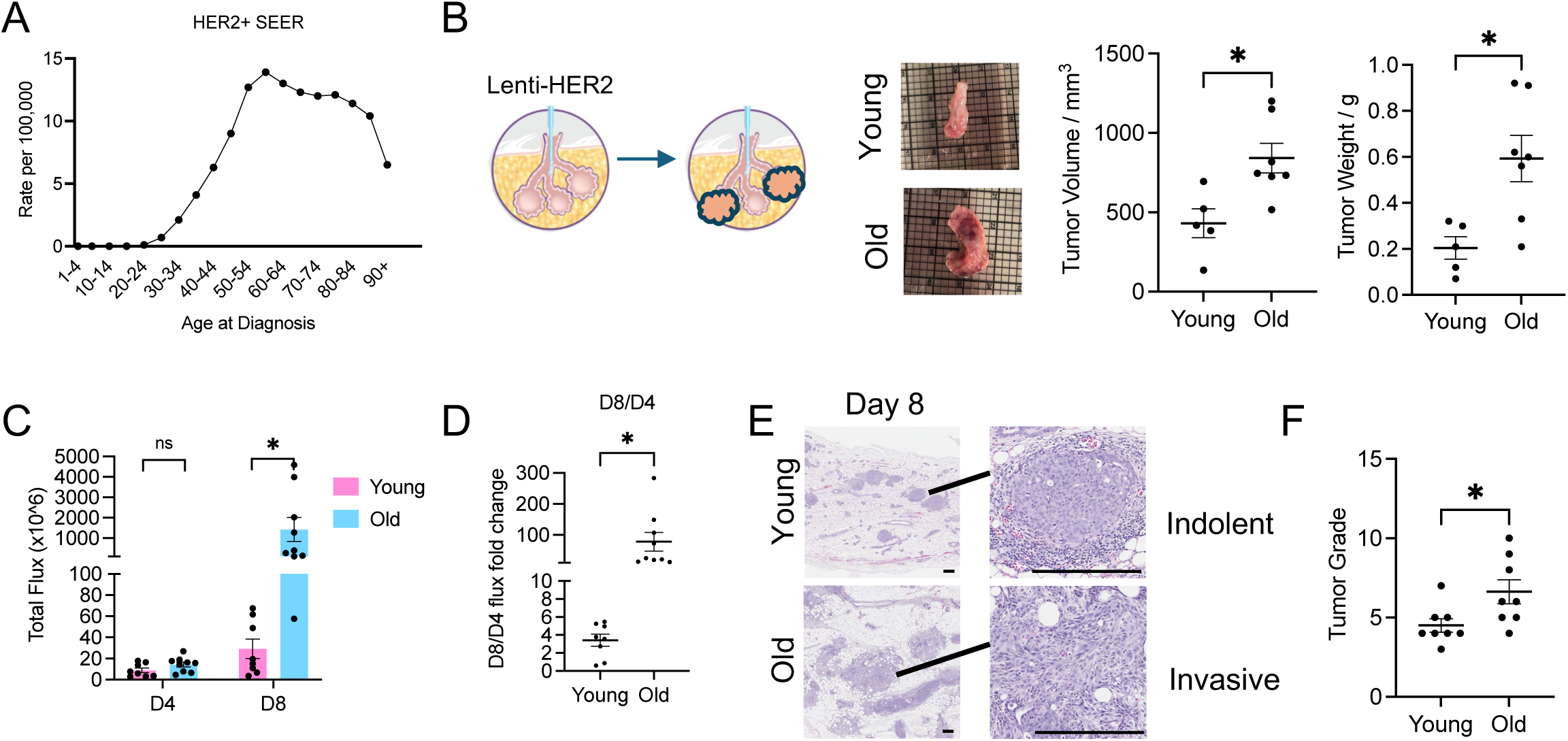
A mouse model of age-dependent breast cancer. (A) Human HER2+ breast cancer incidence with age. (B) The size and weight of breast tumors were measured in mice 14 days after tumor induction. Each dot represents one mouse. Representative tumors shown on the left with the intraductal injection illustration. (C) Total flux of virus injected mammary glands was measured in vivo. (D) The ratio of flux at day 8 and day 4 was calculated. (E) Mammary glands were collected from animals 8 days after intraductal injection and stained by hematoxylin & eosin (H&E) to visualize lesions. Representative lesions are shown as high-power magnification (right). Scale bar: 100 µm. (F) Tumor grade was assessed by Pathology Core at UCSD based on the H&E staining. Unpaired t-test has been used. ns P>=0.05. * P<0.05. *** P<0.001.

Around one week after virus induction, tumors were detected as palpable. On day 14, old tumors were larger than young tumors (Figure 1B). To ensure that the observed difference in tumor burden was not due to variation in virus infection and therefore oncogene transduction efficiency, we co-expressed *HER2*/*neu* with luciferase and measured the total luminescence flux of the injected mammary glands 4 and 8 days after virus injection. On day 4, young and old mice exhibited comparable luciferase expression in successfully injected glands (flux intensity is above 10e6 p/sec/cm2/sr, at least 10 times higher than the background), indicating similar transduction efficiency between young and old mice (Figure 1C, Extended Data Fig. 1A). However, on day 8, luciferase expression was significantly higher in old mice (Figure 1C, Extended Data Fig. 1A). The fold change of flux intensity between day 4 and day 8 was also significantly higher in the old than the young (Figure 1D), suggesting that the age-dependent difference in tumor burden reflects enhanced tumor progression rather than differences in viral infection efficiency.

Histological assessment showed that the development of individual tumor lesions in one gland is not synchronized, and while the majority of the lesions in young mice were indolent with clear membrane boundaries around the tumor, the majority of lesions in old mice were invasive with tumor cells invading through the membrane (Figure 1E). Based on the frequency and severity of invasive lesions, tumor grades were assessed blinded by a pathologist on a scale from 1 to 10. The results showed that, 8 days after tumor induction, the mean tumor grade in old mammary glands was significantly higher than in young mammary glands (Figure 1F). Based on these results, we conclude that the tumor burden induced by the same activated oncogene is greater in old mice than in young mice, and tumors appear histologically more invasive in old mice than young mice.

### Old tumors exhibit gene expression signatures relevant to age-dependent tumor phenotype

We asked if young and old tumors exhibit distinct gene expression signatures. To address this, we applied spatial whole transcriptome profiling by GeoMx and single nucleus RNA-seq (snRNA-seq) to mammary tissues of young and old mice 8 days post tumor induction. For GeoMx, two young and two old mice were compared and five representative tumor lesions were selected as regions of interest (ROI) from one injected mammary gland (mammary gland #4) from each mouse. CD45, panCK, and HER2 antibodies were used to distinguish the immune cell (CD45+), normal epithelium (panCK+, HER2-) and tumor cell (panCK+, HER2+) compartments (Figure 2A). PCA showed the samples clustered primarily by cell compartment, not by age (Extended Data Fig. 2A). Gene expression analysis revealed preferential expression of epithelial marker genes in the HER2+ compartment and expression of immune cell marker genes in the CD45+ compartment, validating the specificity of this approach (Extended Data Fig. 2B).

**Figure 2:**
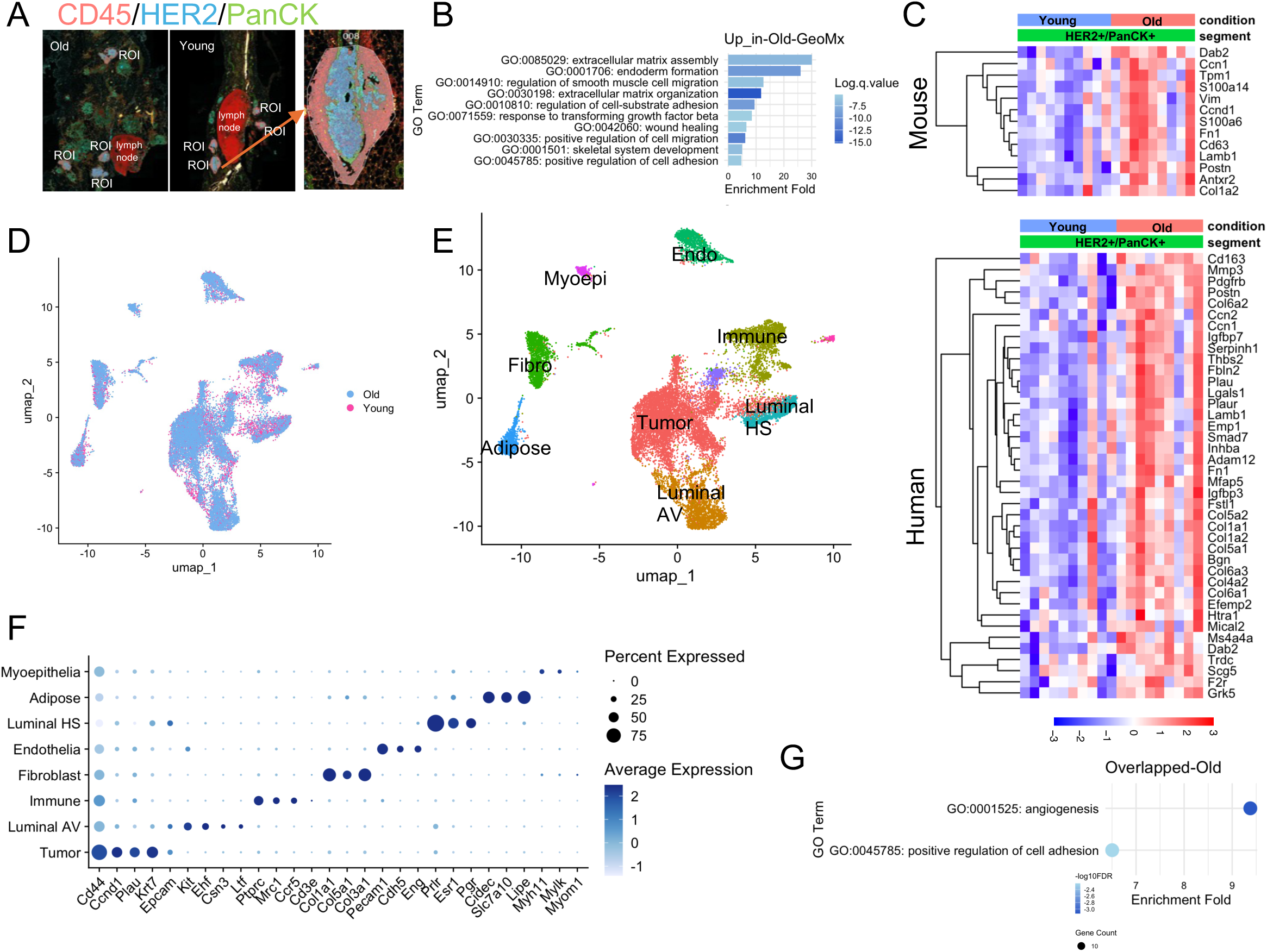
GeoMx and snRNA-seq to investigate transcriptomic changes between young and old mammary tumors. (A) Representative GeoMx image and region of interest (ROI). CD45, HER2, and panCK antibodies mark immune, tumor, and epithelial compartments, respectively. (B) Gene ontology (GO) analysis of biological processes enriched in old tumors based on differentially expressed genes (DEGs) identified from GeoMx tumor compartment (panCK+/HER2+). (C) Expression of selected genes from GeoMx tumor compartment data. Upper panel gene list is from GSEA WANG_TUMOR_INVASIVENESS_UP.v2023.2.Mm. Lower panel gene list is from GSEA SCHUETZ_BREAST_CANCER_DUCTAL_INVASIVE_UP_v2023.2.Hs. (D) UMAP visualization of young and old tumors by snRNA-seq. (E) UMAP showing major cell types identified by snRNA-seq. (F) Expression of canonical marker genes across cell types in snRNA-seq. (G) Overlapping biological processes upregulated in old tumors identified by both GeoMx and snRNA-seq.

From the HER2+/panCK+ tumor compartment, we identified 257 upregulated genes and 91 downregulated genes in the old compared to young (p value < 0.05 and |log2FC| > 0.5) (Supplementary File 1). Gene ontology analysis showed that the top 10 pathways enriched in genes upregulated in the old tumors were these associated with regulation of extracellular matrix assembly, cell migration and cell adhesion (Figure 2B). Indeed, many of the genes upregulated in old tumors are also upregulated in mouse invasive tumors or human invasive ductal breast cancers based on GSEA datasets (Figure 2C), aligning with the more invasive phenotypes of the old tumors (Figure 1E, F).

For snRNA-seq, same as GeoMx, we profiled mammary glands from two young and two old mice collected 8 days after tumor induction. After quality control filtering, 9,237 cells from young mice and 15,444 cells from old mice were retained for downstream analysis. Clustering analysis revealed similar distributions between young and old samples, with no cluster uniquely present in either group (Figure 2D). We identified nine major cell types spanning epithelial and stromal compartments (Figure 2E). Canonical marker genes and markers from prior single cell mammary studies were used to confirm cell identities (Figure 2E, F)^21,22^. Among epithelial populations, we identified tumor cells (expressing *Cd44*, *Ccnd1*, *Plau*, *Krt7*), luminal alveolar (AV) cells (expressing *Kit*, *Ehf*, *Csn3*, *Ltf*), luminal hormone-sensing (HS) cells (expressing *Prlr*, *Esr1*, *Pgr*), and myoepithelial cells (expressing *Myh11*, *Mylk*, *Myome1*). Stromal populations included fibroblasts (expressing collagen genes), adipocytes (expressing *Cidec*, *Lipe*), immune cells (marked by *Ptprc*), and endothelial cells (expressing *Cdh5*, *Eng*).

In the tumor cluster, we identified 936 upregulated and 66 downregulated genes (p.adj < 0.05, |log2FC| > 0.5; Supplementary File 2) in old compared to young. 62 of the 257 genes upregulated in GeoMx were also upregulated in snRNA-seq, representing a highly significant overlap (Fisher’s exact test of overlap between the two gene lists shows overlap is more than expected by chance, p = 4.898e-10). Gene ontology showed that these overlapping genes were enriched in angiogenesis and cell adhesion (Figure 2G), again potentially associated with the more invasive phenotypes in the old tumors. Based on these results, the old tumor cells upregulate gene expression programs that correlate with the more invasive histological phenotypes in the old mice.

### Interferon-simulated genes including CXCL9/10 are downregulated in old tumors

Next we turned to the downregulated genes in the old tumors. Interestingly, within the HER2+ tumor compartment, GeoMx revealed that old tumors downregulated genes related to interferon response, reflected in GO terms such as interferon-beta response, JAK_STAT signal pathway and type II interferon response (Figure 3A). GSEA hallmark analysis also revealed that hallmarks of interferon-gamma response and IL6_JAK_STAT3 pathway were downregulated in old tumor cells (Figure 3B). Notably, two interferon-stimulated genes (ISGs) belonging to the CXCL chemokine family, *Cxcl9* and *Cxcl10*, were the two most downregulated genes in the old tumor compartment (Figure 3C).

**Figure 3:**
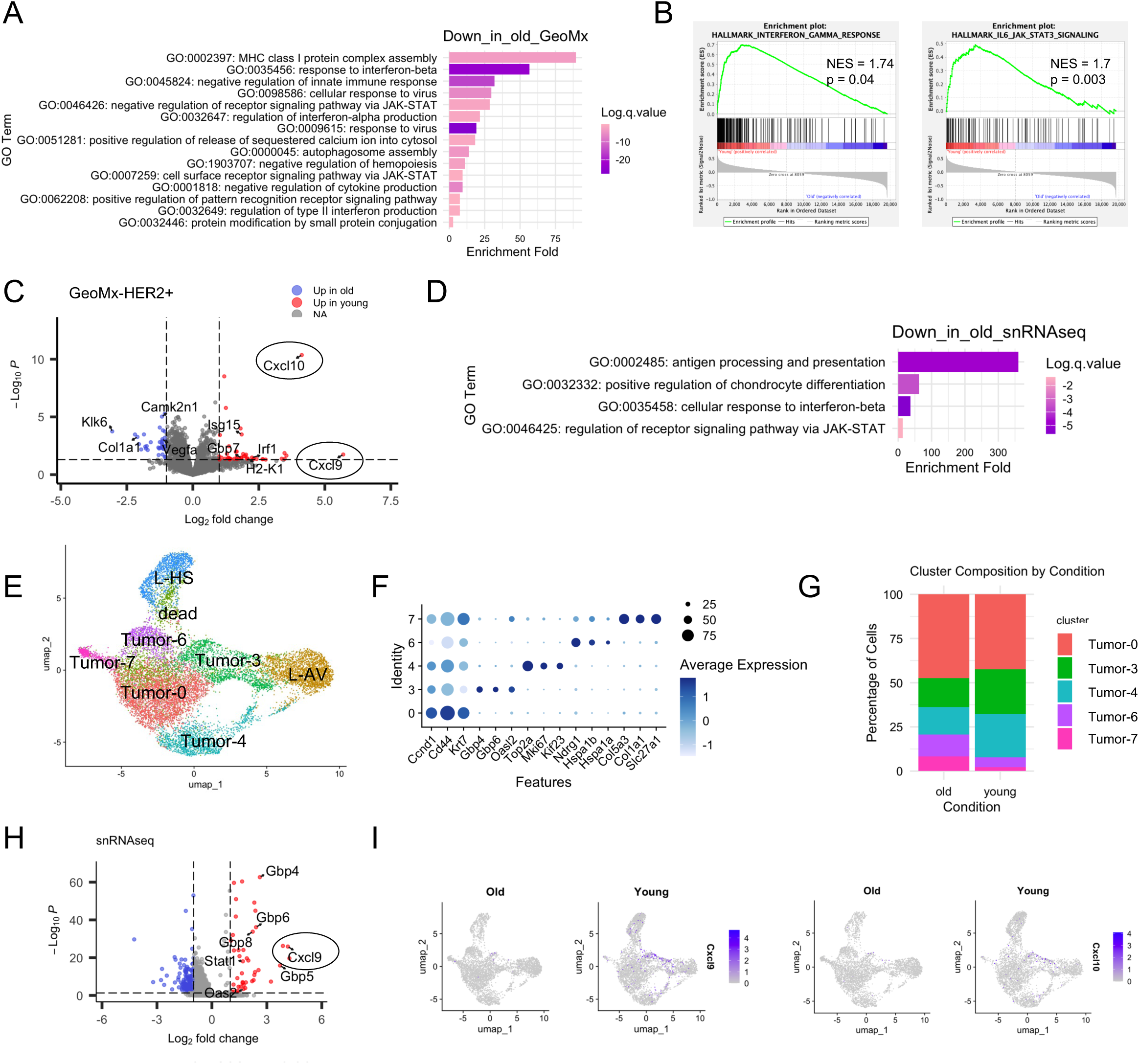
Analysis of downregulated genes in the old tumors by GeoMx and snRNA-seq. (A) GO analysis of biological processes enriched among genes downregulated in old tumor compartments identified by GeoMx. (B) Gene Set Enrichment Analysis (GSEA) of genes downregulated in old tumor compartments from GeoMx. (C) Volcano plot of DEGs from GeoMx tumor compartments, with selected genes highlighted. (D) GO analysis of biological processes enriched among genes downregulated in old tumors identified by snRNA-seq. (E) UMAP visualization of luminal and tumor subclusters identified by snRNA-seq. (F) Expression of representative marker genes across epithelial and tumor subclusters. (G) Proportion of cells in each epithelial/tumor subcluster. (H) Volcano plot of DEGs in subcluster Tumor-3. (I) Feature plots showing *Cxcl9* and *Cxcl10* expression across UMAPs.

Based on GeoMx, the downregulation of *Cxcl9* and *Cxcl10* was apparent in both tumor and immune compartments in old mice, but with a greater fold change observed in the tumor compartment (tumor: *Cxcl10* FC = 4.12, *p* = 4.35 × 10⁻¹¹; *Cxcl9* FC = 5.69, *p* = 0.02; immune: *Cxcl10* FC = 1.87, *p* = 0.08; *Cxcl9* FC = 3.2, *p* = 0.06) (Extended Data Fig. 3A).

Downregulated genes in snRNA-seq Tumor cluster showed strong convergence with GeoMx, with shared enrichment in pathways related to interferon-beta response and JAK-STAT signaling (Figure 3D). GeoMx and snRNA-seq combined suggest a diminished interferon response in aged tumors.

To further investigate the age-associated interferon response in tumor cells, we performed subclustering of Tumor, Luminal AV and HS from snRNA-seq and identified five subclusters within the tumor cells: Tumor-0, 3, 4, 6 and 7 (Figure 3E,F). The Tumor-0 subcluster comprised the majority of cells, with upregulated genes typical of tumor markers, such as *Ccnd1*, *Cd44*, *Krt7* and *Plau*. The Tumor-4 subcluster had proliferating features and was marked by *Mki67* and additional cell cycle genes, such as *Kif23* and *Ndrg1*. The Tumor-3 subcluster was interestingly enriched for ISGs, such as interferon-induced guanylate-binding protein *Gbp* family members and *Oas2*. The Tumor-6 subcluster expressed stress-related heat shock proteins. The Tumor-7 subcluster is a rare population that co-expressed tumor markers alongside extracellular matrix genes. Luminal cells clustered adjacent to the tumor clusters, with luminal AV cells positioned next to the Tumor-3 subcluster, and luminal HS cells loosely connected to Tumor-6 cells (Figure 3E), consistent with luminal cells being the tumor cell of origin.

When comparing the proportions of tumor subclusters between young and old, we observed that young samples had a higher proportion of interferon-related Tumor-3 cells (Figure 3G). When we analyzed the differentially expressed genes in this subcluster, the downregulated genes in the old are enriched in defense response and interferon response pathways (Extended Data Fig. 3B). Several ISGs such as GBP family genes *Gbp4* and *Gbp5*, *Cxcl9*, *Oas2,* and *Stat1* (Figure 3H), were significantly downregulated in the old Tumor-3 subcluster. *Cxcl9* and *Cxcl10* were specifically expressed in the Tumor-3 subcluster in the young, but much lower expressed in old (Figure 3I). From snRNA-seq global clustering, *Cxcl10*+ and Cxcl9+ cells were enriched in a subset of the tumor cluster and an adjacent subset of luminal AV cluster in the young (Extended Data Fig. 3C). The immune cell cluster didn’t show enrichment of *Cxcl9*+ cells (Extended Data Fig. 3C). It is intriguing that *Cxcl9* emerged as one of the most downregulated genes in tumor cells from both GeoMx and snRNA-seq (Figure 3C,H), indicating a potential critical role of *Cxcl9* and *Cxcl10* in the observed phenotypes.

### Validation of CXCL9 and CXCL10 downregulation in old tumors

CXCL9/10 attract CD8⁺ and CD4⁺ T cells and facilitate their recruitment to tumor sites^38,39^. Given this potential functional significance, we next set out to validate the expression of *Cxcl9/10* by RNAscope at the RNA-level and ELISA at the protein level, to confirm that *Cxcl9/10* are downregulated in old tumor cells. By RNA-scope, we confirmed that *Cxcl9* or *Cxcl10* positive cells were more abundant in young tumors and the H-scores of *Cxcl9*/*10* mRNA in young tumors are significantly higher than in old (Figure 4A-D). The *Cxcl9*/*10* positive cells are enriched in the HER2+ tumor compartment, but not the surrounding CD45+ immune compartment. Untransformed mammary ducts from the same slides didn’t express *Cxcl9*/*10*, indicating the expression in young is largely tumor specific (Figure 4C).

**Figure 4.**
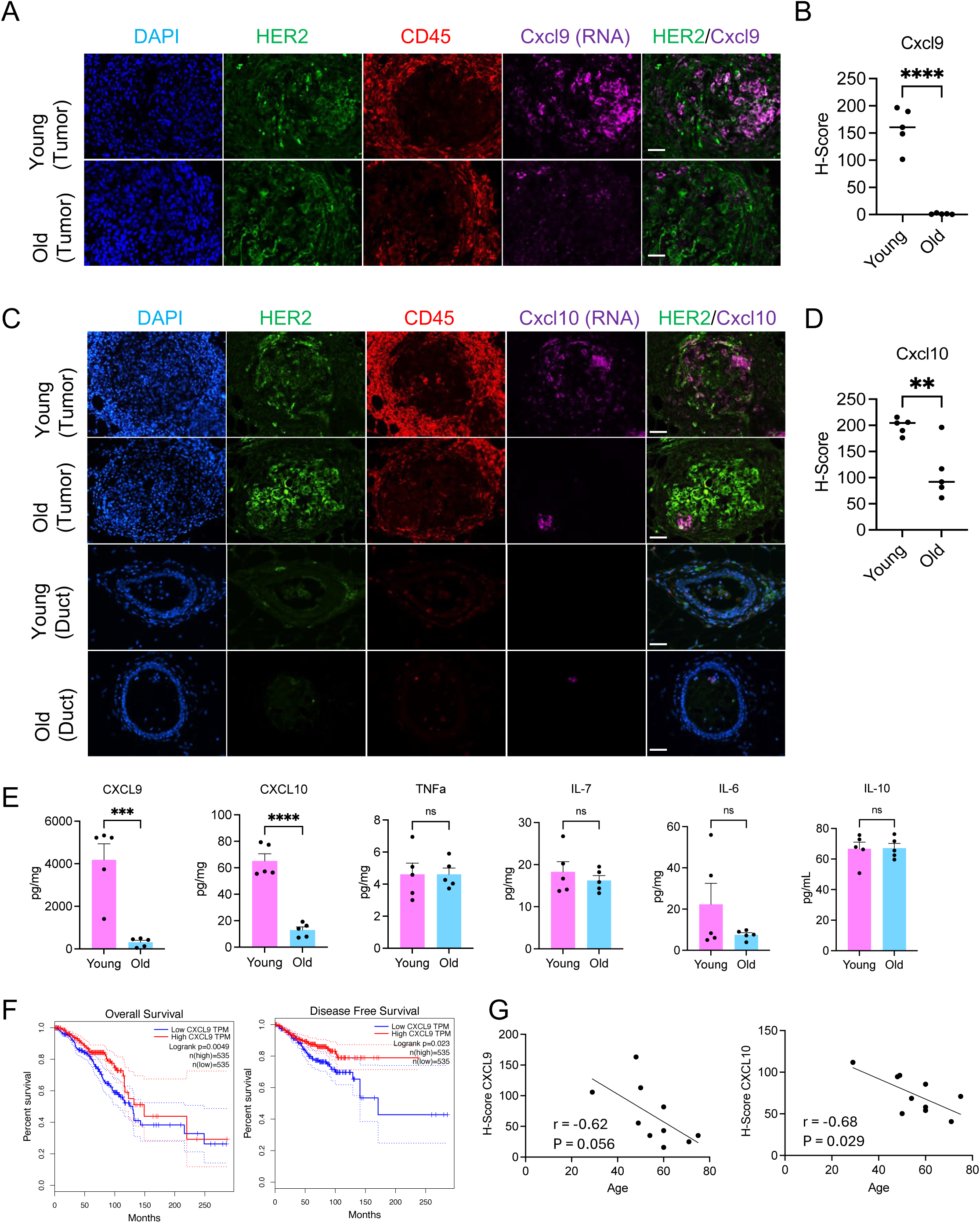
*Cxcl9* and *Cxcl10* expression is reduced in old mammary tumors. (A-D) RNAscope with Cxcl9 (A) or Cxcl10 (C) (purple) RNA probes combined with HER2 (green) and CD45 (red) antibodies in young and old mammary tumors and ducts. (B,D) H-scores were used for statistical analysis on A and C. Each group contained five mice. 5 lesions were analyzed in each mouse. (E) ELISA on mouse mammary tissue homogenate 15 days after the virus injection. (F) The correlation between overall survival and disease-free survival rate and CXCL9 expression from the GEPIA BRCA dataset. (G) Quantification of *CXCL9* and *CXCL10* RNA expression by RNAscope H-score in HER2⁺ patient tumor sections across different ages. Scale bar: 50 µm. For (B,D,E), unpaired t-test was used. ns P >= 0.05. ** P<0.01. *** P<0.001. For (G), Pearson correlation analysis was used.

ELISA on young and old mammary gland whole tissue homogenate two weeks after tumor induction showed that the protein level of CXCL9/10 was also significantly higher in young tumors than in old (Figure 4E). We also tested 30 other commonly studied chemokines such as TNFa, IL family and so on, and none of them showed significant difference between young and old, indicating the specificity of the altered expression of CXCL9/10 (Figure 4E). We conclude that *Cxcl9/10* are specifically downregulated in aged mouse HER2+ tumor cells.

### CXCL9/10 predict better disease outcome and are downregulated in older HER2+ human patients

We asked whether *CXCL9*/*10* expression is correlated with patient outcome and whether this age-dependent downregulation in mouse is also a feature of human tumors. In human TCGA BRCA data^40^, higher *CXCL9* expression level was correlated with better overall survival and disease-free survival in total breast cancers (Figure 4F). *CXCL10* showed a similar trend (Extended Data Fig. 4A). Across all samples, there was no significant correlation between *CXCL9* or *CXCL10* expression and age (Extended Data Fig. 4B). However, using the UCSC Xena browser^41^, in TCGA BRCA samples with *HER2/ERBB2* copy numbers greater than 6, classified as *HER2* amplified tumors^42,43^, we observed a modest trend toward decreased *CXCL9*/*10* expression with increasing age (Extended Data Fig. 4C). To further validate these findings from TCGA data, we performed RNAscope on tissue arrays of human breast invasive ductal carcinoma from patients of varying ages. In the mixed subtype samples, *CXCL9/10* expression showed a modest, non-significant downward trend with age (Extended Data Fig. 4D). In contrast, HER2+ tumors displayed a significant negative correlation between age and the expression of *CXCL9* (r = −0.62, p = 0.056) and CXCL10 (r = −0.68, p = 0.029) (Figure 4G). We conclude that decreased expression of *CXCL9* (and potentially *CXCL10*) is associated with worse patient outcome and downregulation of *CXCL9/10* with age is a feature of human HER2+ breast cancers.

### Downregulation of CXCL9/10 is linked to chromatin accessibility changes

Gene expression changes are often closely linked to alterations in chromatin accessibility. To investigate this relationship globally and at *Cxcl9*/*10* genes in young and old tumor cells, we performed snATAC-seq and snRNA-seq in parallel on the same nucleus preparation. The snATAC-seq data identified the same nine major cell types as snRNA-seq (Figure 5A, compared to Figure 2E). To further investigate the epigenetic landscape in the epithelial and tumor cells, we then subclustered only the tumor and luminal cells, and transferred the subcluster IDs from snRNA-seq in Figure 3E. Among the tumor cells, subclusters Tumor-0 and Tumor-3 were clearly distinguishable, whereas other tumor subclusters identified in snRNA-seq were not as well-resolved in the ATAC-seq data (Figure 5B). Consistent with the snRNA-seq findings (Figure 3G), the interferon-related Tumor-3 subcluster was more prevalent in young samples (27.0%) compared to old samples (15.4%) (Figure 5C).

**Figure 5.**
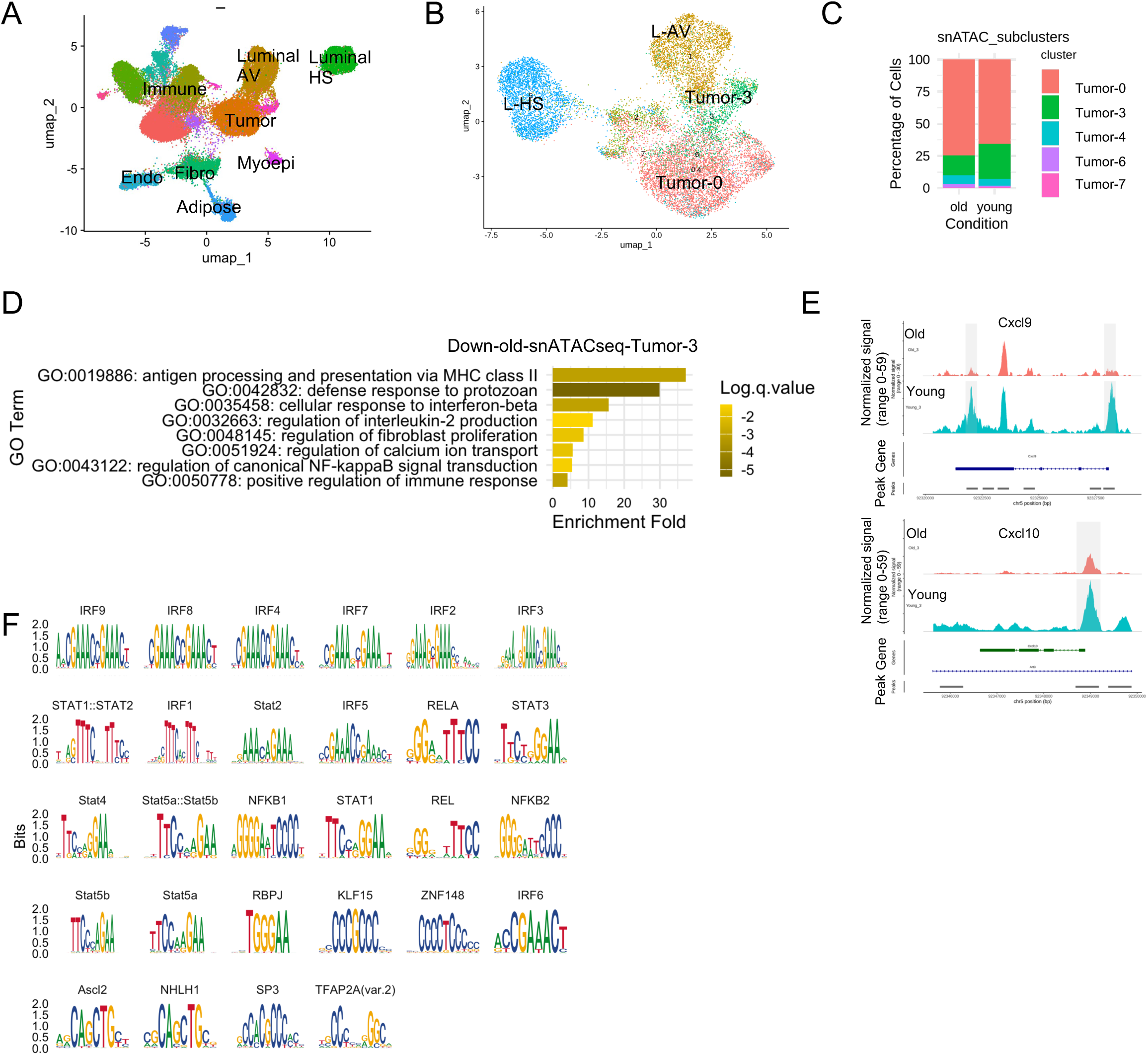
Chromatin accessibility alterations in old tumor cells revealed by snATAC-seq. (A) UMAP visualization of cell types identified by snATAC-seq after annotation transfer from snRNA-seq. (B) Luminal and tumor cell subclustering and annotations transferred from snRNA-seq. (C) Proportion of epithelial subclusters based on (B). (D) GO analysis of genes associated with downregulated chromatin accessibility peaks in old Tumor-3 cells. (E) Chromatin accessibility tracks at the *Cxcl9* and *Cxcl10* loci. Differentially accessible peaks (DAPs) are highlighted in grey. (F) Transcription factor binding motifs enriched in young Tumor-3 accessible DAPs.

We next analyzed differentially accessible peaks (DAPs) within the Tumor-3 subcluster and identified 185 peaks with increased accessibility in young cells and 39 peaks with increased accessibility in old cells (Supplementary File 3). The greater number of accessible peaks in young Tumor-3 cells suggests a generally more open chromatin state. GO analysis revealed that genes associated with DAPs in the young Tumor-3 subcluster were enriched for defense response and interferon response pathways (Figure 5D), similar to Tumor-3 cluster RNA analysis (Extended Data Fig. 3B). Notably, DAPs were localized to the TSS region and the coding sequence near the 3’ end of *Cxcl9*, as well as the TSS region of *Cxcl10* (Figure 5E). In both cases, young samples exhibited markedly higher accessibility signals (Figure 5E), suggesting more open chromatin at *Cxcl9* and *Cxcl10* loci, compared to the closed chromatin state observed in old samples. This is consistent with the increased expression of *Cxcl9*/*10* observed in young tumor cells by GeoMx, snRNA-seq and RNA-scope.

To determine whether the accessible regions in young cells shared common transcription factor (TF) binding motifs and to predict potential TF regulators, we performed motif enrichment analysis on the increased accessible regions in the young Tumor-3 subcluster. We found 28 motifs significantly enriched in these DAPs (padj < 0.05), with interferon regulatory factors (IRFs) and STAT family members, the known drivers of ISGs, being the most prominent (Figure 5F). Notably, IRF1 and STAT1, two established regulators for Cxcl9/10^44,45^, are among the enriched TFs (Figure 5F). These data suggest that old tumors exhibit less accessible chromatin at ISG-associated TF binding motifs, including these associated with *Cxcl9*/*10* genes, and this correlates with decreased expression of *Cxcl9/10* in old tumor cells.

### Tumor-associated T cells are reduced in old tumors

The downregulation of chemokines *Cxcl9*/*10*, which are critical chemokines in T cell recruitment, indicates a dysregulated cancer-immunity cycle in the old tumors^39^. Therefore, we assessed the immune phenotype of the young and old mouse tumors. First, we assessed expression of the T cell marker *Cd3e* in the CD45+ compartment in GeoMx data and found that it was reduced in the old, indicating reduced T cell content (Figure 6A). snRNA-seq is biased toward capturing epithelial rather than immune cells, especially T cells^46,47^. Consistent with this limitation, only 14 *Cd3e*⁺ cells were detected in our dataset. Therefore, we did not place strong emphasis on interpreting the immune cell clusters from the snRNA-seq analysis. However, GeoMx revealed no obvious difference in expression of canonical T cell activation markers (*Ifng*, *Cd69*, *Il2ra*/*Cd25*, *Gzmb*, etc.) and exhaustion/inhibition markers (*Pdcd1*, *Tox*, *Ctla4*, *Tigit*, *Lag3*, etc.) within the CD45⁺ compartment of young and old mice (Figure 6B).

**Figure 6.**
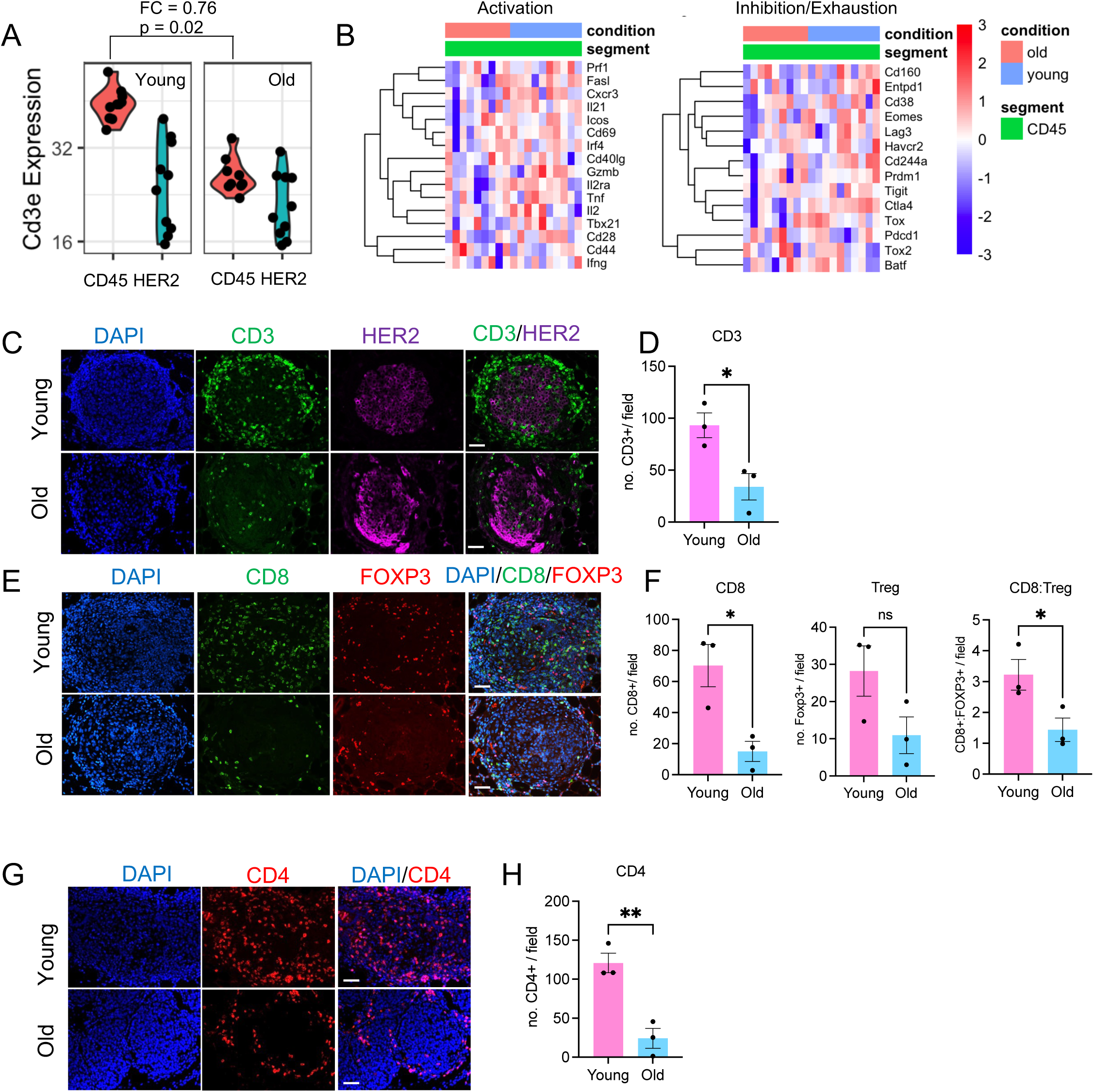
Tumor associated T cells are reduced in old mammary tumors. (A) *Cd3e* expression measured by GeoMx. Each dot represents one ROI. CD45+ and HER2+ compartments are shown. Fold change and p-value for CD45+ compartment from GeoMx analysis are shown. (B) Heatmap of canonical T-cell activation and exhaustion markers in CD45⁺ compartments from GeoMx. (C) Immunostaining of CD3 (green) and HER2 (purple) on mammary tumors. (D) Number of CD3 positive cells per field. (E) Immunostaining of CD8 (green) and FOXP3 (red). (F) Number of CD8 positive cells, FOXP positive cells, and the ratio of CD8:FOXP3 were calculated. (G) Immunostaining of CD4 (red) on mammary tumors. (H) Number of CD4 positive cells per field. For (D,F,H), each dot represents the average number of at least five fields from one mouse. Unpaired t-test was used. ns P >= 0.05. * P<0.05. ** P<0.01. Scale bar: 50 µm.

Aligned with *Cd3e* reduction from GeoMx, immunostaining showed fewer total CD3⁺ T cells in both peri-tumoral and intra-tumoral regions in old mice (Figure 6C-D). Both CD4⁺ and CD8⁺ T cell numbers were diminished in old tumor lesions (Figure 6E-H). Although immune suppressive FOXP3⁺ regulatory T cells (Tregs) showed a trend of increase in young tumors, the CD8⁺/FOXP3⁺ ratio remained significantly higher in the young group (Figure 6E-F). To assess for functional differences in tumor-associated T cells between young and old mice, we isolated CD3⁺ T cells from young and old tumors 15 days post-induction and stimulated them with PMA and ionomycin *in vitro*. Following stimulation, the percentage of CD8⁺ T cells expressing IFNγ or CD44 was similar in young and old mice (Extended Data Fig. 6A), suggesting that tumor-associated T cells from both age groups can be activated at similar levels. Regardless, based on numbers of infiltrating cytotoxic T cells and their balance with immune suppressive Tregs, the immune microenvironment appeared more cytotoxic and anti-tumor in young tumors.

### Downregulation of CXCL9/10 – T cell signaling drives increased tumor burden in old

To test whether increased T cells might be causally involved in tumor suppression in young mice, we depleted CD4⁺ or CD8⁺ T cells in young mice. In young mice, tumor volume significantly increased following depletion of CD4⁺ or CD8⁺ T cells (Extended Data Fig. 7A). Combined depletion of CD4⁺ and CD8⁺ T cells enhanced tumor growth in young mice, but had no significant effect in old mice (Figure 7A). These data suggest that reduction of T cells contributes to the enhanced tumor growth and disease burden observed in old mice, consistent with a causal role of T cells in anti-tumor immunity in young.

**Figure 7.**
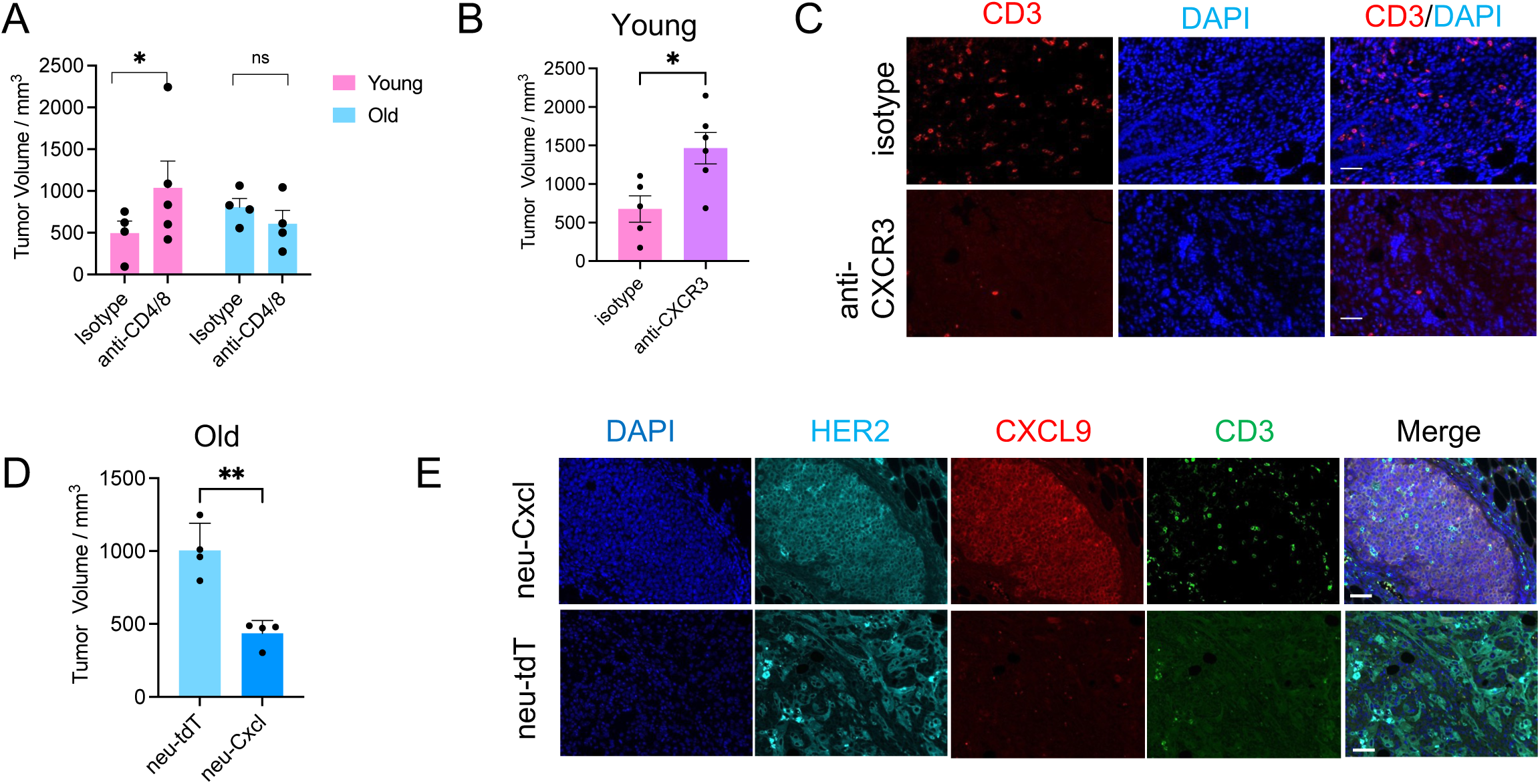
*Cxcl9* and *Cxcl10* are associated with age-dependent HER2⁺ tumor growth. (A)Tumor volumes in mice treated with isotype or combined anti-CD4 and anti-CD8 antibodies were measured 15 days after tumor induction. (B) Tumor volumes measured 15 days after tumor induction in mice treated with isotype control or anti-CXCR3 antibodies. (C) Immunofluorescence staining of CD3 (red) in tumors from (B). (D) Tumor volumes measured 15 days after intraductal injection of lenti-HER2-tdTomato or lenti-HER2–Cxcl9–Cxcl10. (E) Immunofluorescence staining of tumors from (D) showing HER2 (cyan), CXCL9 (red), and CD3 (green). Scale bar: 50 µm. Unpaired t-test was used for statistical analysis. ns P >= 0.05. * P<0.05. ** P<0.01

We then tested the casual role of *Cxcl9/10* downregulation in tumorigenesis. CXCL9/10 recruit T cells by binding the CXCR3 receptor on T cells^38^. To assess whether blocking CXCL9/10 signaling accelerates growth of young tumors, we inhibited CXCR3 using a neutralizing antibody. FACS confirmed that the CXCR3+ cells were decreased from ∼25% to ∼5% of total CD3 following CXCR3 antibody treatment (Extended Data Fig. 7B,C). Two weeks after tumor induction, young mice treated with CXCR3 blockade developed larger tumors (Figure 7B) and exhibited reduced tumor-associated T cell infiltration (Figure 7C).

Conversely, to test whether enhancing *Cxcl9/10* expression could suppress tumor growth in old mice, we co-expressed mouse *Cxcl9/10* with the same *Her2/neu* used in this study using a lentivirus P2A expression system. Tumor grade was similar between Cxcl9/10- and tdTomato-overexpressing groups, suggesting that these chemokines do not affect tumor invasiveness (Extended Data Fig. 7D). However, compared to Her2-tdTomato controls, Her2-Cxcl9-Cxcl10 infected mammary glands showed increased tumor-associated T cell infiltration and reduced tumor burden (Figure 7D,E). In sum, we conclude the suppression of *Cxcl9* and *Cxcl10* in old tumors drives decreased infiltration of anti-tumor T cells and faster growth of the tumor.

## Discussion

This study has leveraged a novel mouse model of age-dependent breast cancer to unmask impaired crosstalk between aged tumor cells and immune cells that promotes tumor progression in old. More specifically, our study demonstrates that aged human and mouse breast tumor cells produce lower levels of the T cell–attracting chemokines CXCL9 and CXCL10. This reduction contributes to an immunosuppressive tumor microenvironment and promotes tumor growth. These findings highlight a causal connection between age-associated defects in immune activation by tumor cells and accelerated tumor development.

We employed intraductal injection of the rat oncogene *HER2/neu* into mouse mammary glands to induce breast tumors. This approach offers distinct advantages: while the process is initiated by oncogene activation in normal epithelial cells, it allows us to examine the pro-tumorigenic effects of aging on both epithelial-derived tumor cells and the mammary microenvironment. We selected the HER2 oncogene because incidence of HER2+ breast cancer increases with age in humans (Figure 1A) and delivering HER2 via lentivirus has been successfully used to model breast cancer in young mice^34–36^, and because it represents a clinically relevant subtype not driven by hormonal changes. This is important as aged women undergo profound hormonal shifts that are not paralleled in aged mice^48^.

We reported that CXCL9/10 mediated T cell recruitment drives the age-dependent phenotypes in model. However, systematic decline of T cell function with age also contributes to the enhanced tumorigenesis in older individuals^5,33^. Therefore, we suspect that suppression of CXCL9/10 is not the only mechanism for the enhanced tumorigenesis in old. Future single-cell RNA-seq profiling of CD45⁺-sorted populations may provide deeper insight into age-associated alterations in the immune microenvironment.

CXCL9 and CXCL10 can be secreted by tumor cells and immune cells^38,45,49^. In our model, CXCL9/10 expression was detected in both tumor and immune cells, with tumor cells being the predominant source, as supported by GeoMx, RNAscope, and snRNA-seq analyses. However, the underrepresentation of immune cells in the snRNA-seq dataset may have limited our ability to fully characterize CXCL9/10-producing immune subsets at the single-cell level. Notably, a subset of luminal cells clustering adjacent to tumor cells by UMAP showed enrichment of Cxcl9 expressing cells (Extended Data Fig. S3C). However, comparison of healthy mammary glands from young and old mice revealed no age-related differences in expression and chromatin accessibility at the *Cxcl9* and *Cxcl10* loci (RNAscope and data not shown). These data combined suggest that transcriptional alteration and epigenetic remodeling of *Cxcl9/10* genes occur upon oncogene activation, and that a subset of luminal cells in oncogene-infected glands exhibit both luminal and tumor-like transcriptional features, which might be oncogene-expressing pre-malignant cells.

In the clinic, younger breast cancer patients are often considered to present with more aggressive tumors, largely because more aggressive subtypes such as HER2-positive and triple-negative disease are proportionally enriched in younger cohorts^50^. Importantly, however, older patients experience higher mortality across all subtypes, primarily due to reduced tolerance of standard therapies^16–18^. Our results suggest that boosting chemokine production, particularly CXCL9 and CXCL10, in combination with first-line treatments, may improve outcomes in older patients. In fact, many studies have demonstrated potential strategies to enhance CXCL9/10 expression to enhance anti-tumor immunity: intratumoral delivery of a TLR2 agonist (FSL-1) with IFN-γ was shown to enhance CXCL10 expression and T cell infiltration in a TNBC mouse model^51^, and LNP-delivered IFN-α2 suppressed lung metastases through upregulation of Cxcl9^52^. Clinical trial NCT04830592 uses intravenous infusion to deliver an adenoviral vector producing a bispecific T cell activator (TAc) plus CXCL9/CXCL10/IFNa2 to kill tumor cells and stimulate immunity against the tumor cells^53^. Such approaches merit investigation as therapeutic avenues for older HER2-positive breast cancer patients, a population that is typically under-represented in clinical trials.

Our study underscores the importance of epigenetic regulation in both aging and tumorigenesis. We observed that age-related differences in gene expression in tumor cells are associated with altered chromatin accessibility (Figure 5E-G). Future work dissecting how additional epigenetic mechanisms might regulate age-associated *Cxcl9*/*10* expression changes in tumor cells, such as DNA methylation and histone modifications^45,54,55^, will be critical to fully understand how aging shapes the cancer epigenome.

A limitation of our approach is that virus-mediated oncogene delivery to the epithelium *in situ* likely infects multiple or many cells, thereby not faithfully recapitulating the very rare stochastic mutation followed by clonal expansion that is thought to give rise to human cancer^56^. However, in histopathological analysis, at early stages of disease we do observe discrete well-separated neoplastic lesions (Figure 1E), suggesting that in this model tumors do arise from clonal expansion of isolated single cells. Thus, we think this is a valuable model that complements transplant models of age-dependent breast cancer.

Another limitation of our approach is that it is challenging to know whether the absence of tumors (Supp. Fig 1A) results from failed intraductal injection of virus or effective anti-tumor immune defense. Given this limitation, our study focused on tumor progression, and only mice that developed tumors, which either have higher than 10^6 photon/sec flux with luciferase or harbor visible tumors after dissection with tdtomato, were included in the analyses.

We also note that age-dependent tumor phenotypes are highly model dependent. For example, in a recent lung cancer study, aging was reported to suppress tumor growth^57^. In a HER2⁺ cell line transplantation model, tumors also grew more rapidly in young mice^28^. In contrast, TNBC cell line from the same study showed that it grew faster in old mice^28^. Other studies also show that aging promotes tumor growth^25,26,33^. Factors such as mouse age, genetic background, and the method of tumor induction can all contribute to these differences. These divergent outcomes underscore the complexity of modeling age-related effects in cancer and suggest that each mouse model captures distinct aspects of the human disease. In this case, we showed that downregulation of CXCL9/10 is a feature of both mouse and human HER2+ breast tumors, supporting the tumor-promoting role of aging.

In conclusion, our study reveals that aging profoundly shapes the crosstalk between mammary tumor cells and microenvironment, diminishing tumor cell–driven immune activation through reduced chemokine expression and thereby promoting tumor progression. These findings underscore aging as a mechanistic determinant of breast cancer incidence and outcomes. By establishing an in situ model that captures age-related differences in both tumor cells and their microenvironment, we provide a powerful framework for mechanistic and therapeutic discovery. Ultimately, this work highlights novel opportunities to improve outcomes for older breast cancer patients by restoring immune surveillance and targeting age-specific vulnerabilities.

## Methods

### Animals

This study was approved by the Institutional Animal Care and Use Committee at Sanford Burnham Prebys MDI (AUF 22-032, 23-052). Aged BALB/c-ByJ-NIA mice were requested from NIA. Young BALB/c-ByJ mice were purchased from Jackson Labs and group housed at 21–24 °C, 30–70% humidity, and a 12 h light/dark cycle under specific pathogen free conditions with ad libitum access to water and food.

### Breast tumor induction and measurement

To induce tumor in mice, lentivirus carrying truncated rat *HER2/neu(Erbb2)*-Luciferase or *HER2/neu*-Tdtomato were injected into mouse mammary 4th left glands. Mice were anesthetized via isopropanol inhaling during injection and then put back to cages right after injection. 10^5 TU virus was injected for each gland.

For tumor volume and weight measurement, mice were euthanized under CO2 and the mammary glands with virus injection were dissected out. The glands were weighted and measured width, length and height. Weight x length x height was calculated as volume. Only mice with tumors were included in the analysis.

For luminescence flux measurement, 150 mg/kg luciferin was i.p. injected into each mouse, anesthetized with isoflurane gas, and waited 10 minutes before imaged using IVIS system. Total flux (photons/second) for same size of ROIs was measured.

### In-vivo blockade

Antibodies were prepared in 100 μL sterile PBS and intraperitoneally injected into mice 2 days before the tumor induction, and then once per week afterwards till the end of the experiment. 100 μg antibodies per mice was used. The depletion efficiency was measured by FACS one week post antibody injection.

Antibodies used for injection: CD8b (Bioxcell. BE0223), CD4 (Bioxcell. BE0003), CXCR3 (Bioxcell. BE0249), Rat IgG1 isotype control (Bioxcell. BE0088).

Antibodies used for FACS: CD8a (BioLegend, Cat# 100726), CD8b (BioLegend, 140408), CD4 (BioLegend, 100432), CD3 (Biolegned, 100204), CXCR3 (BioLegend, 126515).

### H&E staining

Hematoxylin and eosin (H&E) staining was performed by the Histology Core at SBP. H&E stained slides were scanned using the Aperio AT2 slide scanner (Leica Biosystems), and presented using Aperio ImageScope (Leica Biosystems). Tumor grade was accessed by pathologist by randomly choose 5 regions on a slide.

### Immunostaining

Immunofluorescence staining was performed on paraffin-embedded sections following standard protocols. Briefly, sections were deparaffinized, rehydrated, and subjected to heat-induced antigen retrieval in citrate buffer (10 mM sodium citrate, pH 6.0) or Tris-EDTA buffer (pH 9.0), depending on the target antigen. Sections were blocked in 5% BSA containing 0.1% Triton X-100 for 30 mins at room temperature and then incubated overnight at 4 °C with primary antibodies. After washing, sections were incubated with species-appropriate Alexa Fluor-conjugated secondary antibodies (Invitrogen) for 1 hour at room temperature. Nuclei were counterstained with DAPI, and slides were mounted. Images were captured using Nikon T2 microscope and processed with ImageJ.

Immunostaining: HER2/ERBB2 (Santa Cruz Biotechnology, sc-33684). CD3 (Abcam. ab16669). CD4 (Abcam. ab183685). CD8 (Abcam. ab217344). CD45 (Abcam. ab10558). CXCL9 (R&D. AF492SP)

### RNAscope

RNAscope in situ hybridization combined with immunofluorescence was performed on formalin-fixed paraffin-embedded (FFPE) tissue sections using RNAscope® Multiplex Fluorescent v2 kit following the protocol RNAscope® Multiplex Fluorescent v2 Assay combined with Immunofluorescence - Integrated Co-Detection Workflow (ICW) (Advanced Cell Diagnostics, ACD). Images were acquired using imaged on a Nikon T2 microscope. H-score was analyzed by software Qupath to semi-quantify the percentage of probe dots following the protocol Using QuPath to analyze RNAscope™, BaseScope™ and miRNAscope™ images from ACD.

Probes were ordered from ACD: human Cxcl10 (cat: 311851), human Cxcl9 (cat: 440161-C2). Mouse Cxcl9 mouse (cat: 489341-C3). Mouse Cxcl10 (cat: 408921). Human patient microarray slides were purchased from Tissuearray.com. Slides Her2d and BR042B were used in the analysis.

### GeoMx Sample Preparation, Sequencing, and Data Analysis

Tissue samples were prepared using GeoMx Digital Spatial Profiler (Nanostring Technologies, Seattle, WA) following the manufacturer’s guidelines. FFPE tissue sections were deparaffinized and rehydrated using standard xylene and alcohol treatments. Sections were then incubated with mouse whole transcriptome atlas panel.

After hybridization, tissues were washed and stained with antibodies and syto13 for nuclear counterstaining. Sections were subjected to GeoMx Digital Spatial Profiler system for regions of interest selection. Probes were captured by laser and collected for sequencing. Raw sequencing data were processed by NanoString GeoMx NGS Pipeline using GUI on a windows computer locally following manufacturer’s guideline. Processed DCC files were then analyzed using R GeoMxTool pipeline (https://www.bioconductor.org/packages/release/workflows/vignettes/GeoMxWorkflows/inst/doc/GeomxTools_RNA-NGS_Analysis.html). EnhancedVolcano package was used to generate volcano plot.

### Single-nucleus RNA sequencing and analysis

Mammary glands were dissected from mice 8 days after tumor induction and snap frozen using liquid nitrogen. Frozen tissues were processed for nuclei isolation using the Nuclei Isolation Kit (Miltenyi Biotec) in combination with the gentleMACS Dissociator (Miltenyi Biotec), according to the manufacturer’s protocol. Nuclei concentration and viability were assessed using an automated cell counter (K2 Cellometer, Nexcelom). Single nuclei capture and library preparation were performed with the Chromium X platform (10x Genomics) and Chromium Next GEM Single Cell 3′ Reagent Kit v3.1 (10x Genomics), targeting 10,000 cells per sample. Two batches of young and old samples were processed. Tissues were processed the same but sequenced on different platforms. For the first batch (one young and one old), sequencing was performed on an Element Biosciences AVITI instrument with the Cloudbreak FS High Output kit, for the second batch (one young and one old), sequencing was Illumina NextSeq 2000.

Raw sequencing data were processed with cellranger 8.0.0 using the mm10 reference genome. Seurat R package was used for downstream analysis. Low-quality nuclei were excluded. Counts were log-normalized and scaled to 10,000 transcripts per nucleus. The package DoubletFinder was used to exclude cells likely to be doublets.

Data were scaled to regress out technical factors such as total UMI counts and mitochondrial gene percentage. Principal component analysis (PCA) was performed, and nuclei were clustered using a shared nearest neighbor (SNN) graph and the Louvain algorithm. Uniform Manifold Approximation and Projection (UMAP) was used for dimensionality reduction and visualization. Marker genes for each cluster were identified with Seurat’s FindMarkers function using the Wilcoxon rank-sum test. DEG analysis between conditions was performed using FindMarkers function.

### Single-nucleus ATAC sequencing and analysis

For snATAC-seq, same nuclei extracted for RNAseq was used and processed by Chromium Single Cell ATAC kit (10x Genomics). Raw sequencing data were processed with cellranger-arc 2.0.2 using the mm10 reference genome. The resulting fragment files and peak-barcode matrices were imported into the Signac and Seurat R packages for downstream analysis. Nuclei were filtered based on quality-control metrics. The peak count matrix was normalized, and dimensionality reduced using SVD on the top features. Low-dimensional embeddings were generated with Uniform Manifold Approximation and Projection (UMAP). Clustering was performed using a shared nearest neighbor (SNN) graph constructed from the reduced dimensions.

Differentially accessible chromatin regions were identified between conditions using Signac’s FindMarkers function. Peaks were annotated to nearby genes using ChIPseeker. Motif enrichment analysis was performed using the FindMotifs function in Signac with the JASPAR 2020 motif database.

To integrate transcriptional and chromatin accessibility data, the Seurat/Signac multiome workflow was used. First, gene activity scores were computed for each nucleus in the snATAC-seq dataset by aggregating accessibility signals across gene bodies and promoter regions. The gene activity matrix was normalized and scaled in the same manner as the snRNA-seq expression matrix. Integration anchors were identified between the snRNA-seq expression matrix and the snATAC-seq gene activity matrix using Seurat’s FindTransferAnchors function. The anchors were then used to transfer cell-type labels and to jointly visualize RNA and ATAC data in a shared UMAP embedding.

### Gene ontology analysis

Gene ontology on the DEGs and DAPs was analyzed by Metascape online tool^58^ (http://metascape.org) and visualized by R ggplot2. q.value less than 0.05 was set as a cutoff except Figure 2A, where p.value 0.05 was used as a cutoff.

### T cell in vitro activation

Mammary glands were dissected on the day of the experiment and dissociated into single cells using the Mouse Tumor Dissociation Kit (Miltenyi, 130-096-730) and the gentleMACS Octo Dissociator soft tissue program. Following dissociation, cells were washed with PBS, treated with ammonium chloride buffer for 2 min to lyse red blood cells, and resuspended in PBS containing 5% BSA. T cells were enriched using the Mouse CD3 T Cell Isolation Kit (BioLegend, 480023) and counted using trypan blue and a hemocytometer.

Enriched T cells were seeded into 12-well plates and stimulated with Cell Activation Cocktail with Brefeldin A (BioLegend, 423303) for 4 h at 37 °C. Cells were then stained with LIVE/DEAD Fixable Dye (Invitrogen, Cat# 65-0865-14) for 15 min at room temperature, pelleted, and incubated with anti-CD16/32 (1:500; TruStain FcX™ PLUS, BioLegend, 56603) to block Fc-receptor binding. Surface staining was performed for 30 min at 4 °C using antibodies against CD45 (BioLegend, 103125), CD8a (BioLegend, 100726), and CD44 (BD, 741057).

Intracellular staining was carried out using the FoxP3 Transcription Factor Staining Kit (eBioscience). Anti-IFNγ antibody (BioLegend, 505809) was diluted in 1× permeabilization buffer and incubated with cells for 45 min at 4 °C. After staining, cells were fixed in 2% paraformaldehyde, resuspended in 100 µL 1× PBS, and acquired within 24 h on a Cytek Aurora 5L spectral flow cytometer. Data were unmixed using SpectroFlo (Cytek Biosciences) and analyzed in FlowJo to generate representative plots and quantification.

### Statistical Analysis

Statistical analysis methods are described in the figure legends.

## Data Availability

GeoMx, snRNA-seq and snATAC-seq data will be available on Gene Expression Omnibus.

## Acknowledgements

This work was supported by P01 AG073084 (PDA), U54 AG079758 (PDA), R01 AG071861 (PDA), NCI Core Grant P30 CA030199-44 (PDA), NCI R01CA174869 (JY), R01CA268179 (JY), Curebound Discovery Grant 23DG10 (JY), Krueger v. Wyeth research award (JY), and San Diego Nathan Shock Center Pilot Grant 2023 (SY).

We also thank the supports from SBP Virus Core (P30 CA030199 and Shared Instrumentation Grant S10 OD036254), SBP Flow Cytometry Core (NCI CCSG P30CA030199, Shared Instrumentation Grant S10OD032325), SBP Genomics Core, SBP Histology Core, SBP Animal Facility and NGS Core from Salk institute (NIH-NCI CCSG P30 CA014195, NIH-NIA San Diego Nathan Shock Center P30 AG068635).

## Author contributions

Conceptualization: S.Y, K.T.Y, J.Y, P.D.A. Data generation and analysis: S.Y, K.T.Y, K.L, A.G, A.D, R.A. Computation: S.Y, S.M, X.L. Writing: S.Y, J.Y, P.D.A. Funding: S.Y, P.D.A. Supervision: P.D.A.

## Extended Data Figures

**Extended Data Fig. 1.**
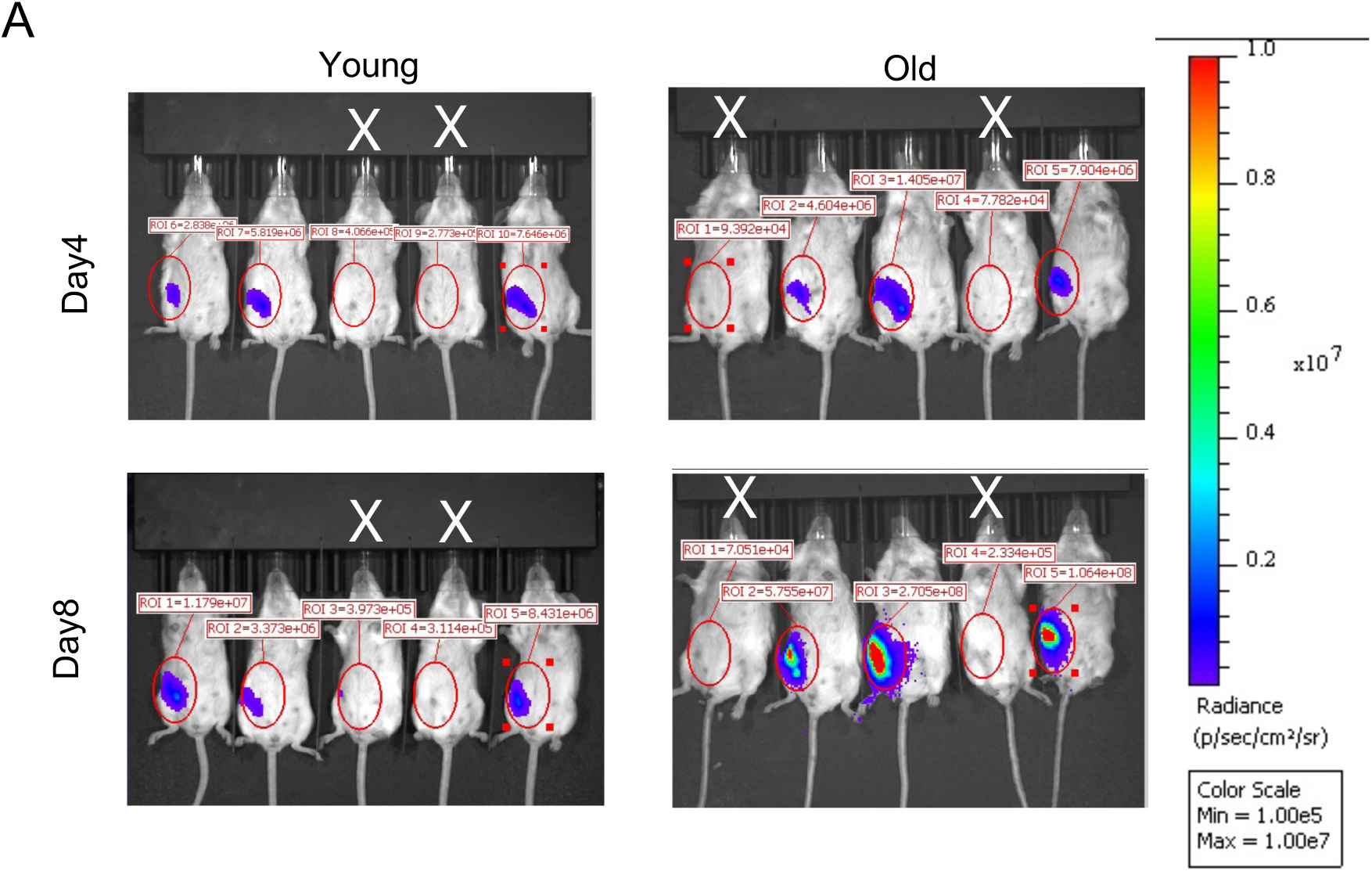
IVIS of mice injected with lenti-HER2-luciferase. Related to Fig. 1. (A) Representative IVIS images of mice following lenti-HER2-Luc injection. Mice marked with a white “×” indicate failed injections and were excluded from analysis. The flux intensity scale bar is shown on the right.

**Extended Data Fig. 2.**
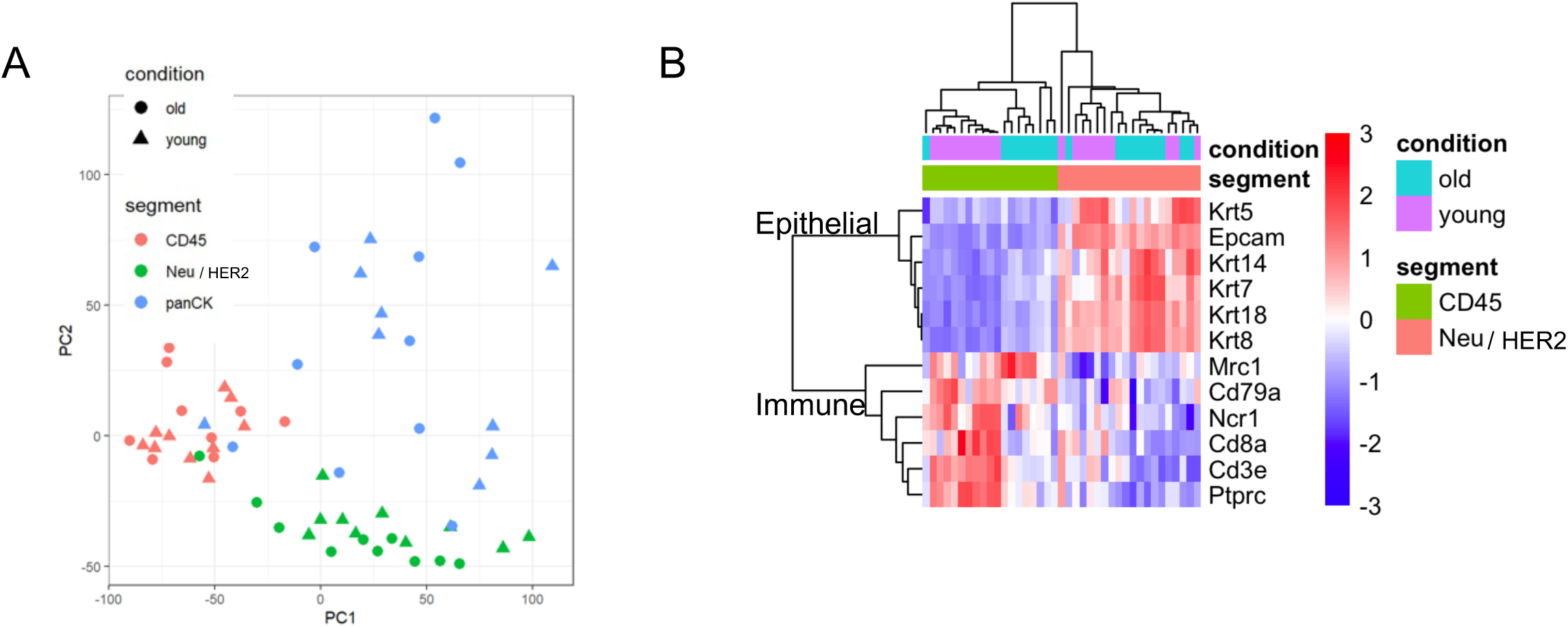
PCA and cell type marker expression from GeoMx. Related to Figure 2. (A) PCA of the GeoMx ROIs. (B) Heatmap showing the expression of epithelial cell markers and immune cell markers in CD45+ and Neu/HER2+ cell compartments.

**Extended Data Fig. 3.**
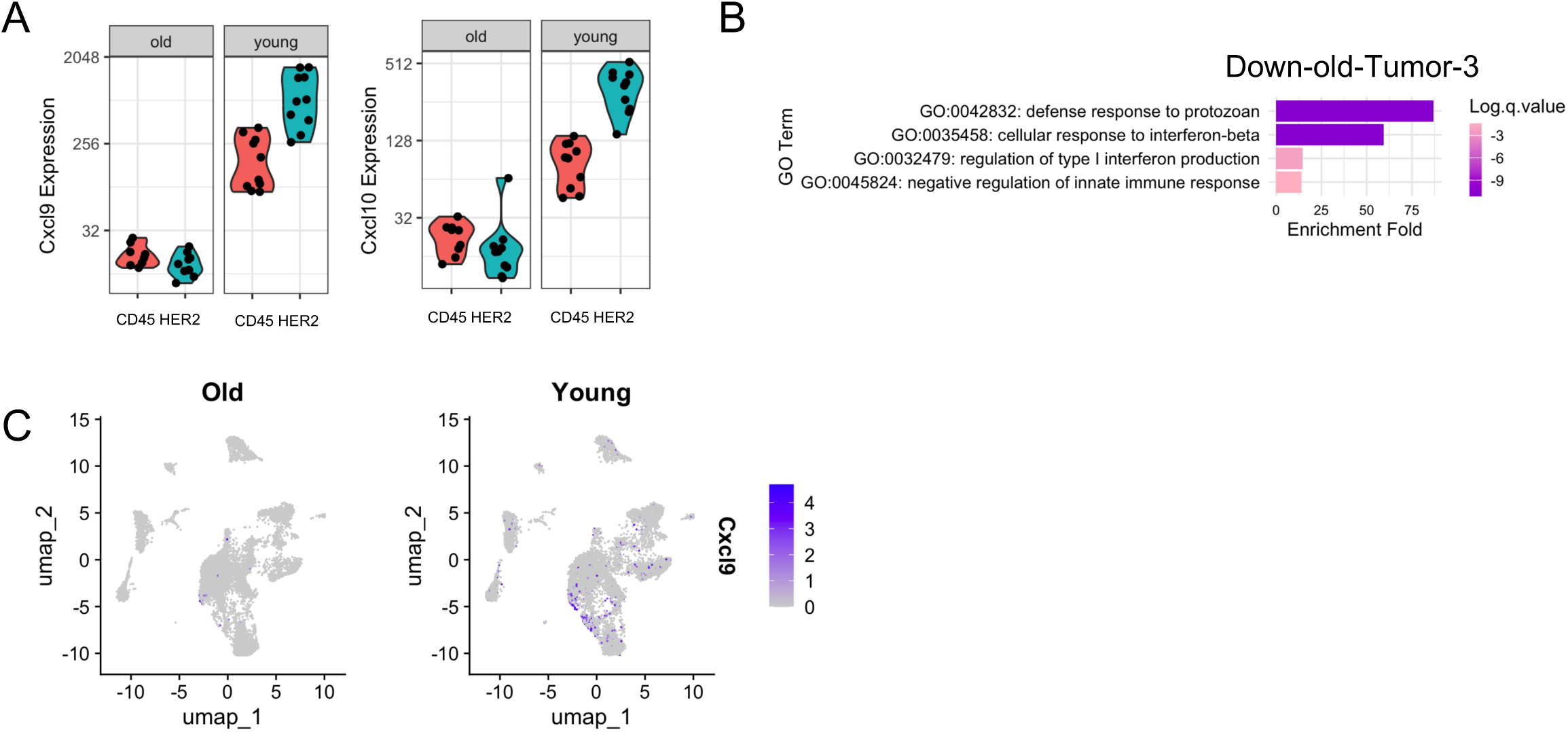
Expression of Cxcl9 and Cxcl10 from GeoMx and snRNA-seq. Related to Figure 3. (A) Expression of *Cxcl9* and *Cxcl10* measured by GeoMx. Each dot represents one ROI. CD45+ and HER2+ compartments were shown. (B) GO analysis of biological processes enriched among genes downregulated in old subcluster Tumor-3 in snRNA-seq. (C) *Cxcl9* positive cells distribution across cell types on the snRNA-seq UMAP.

**Extended Data Fig. 4.**
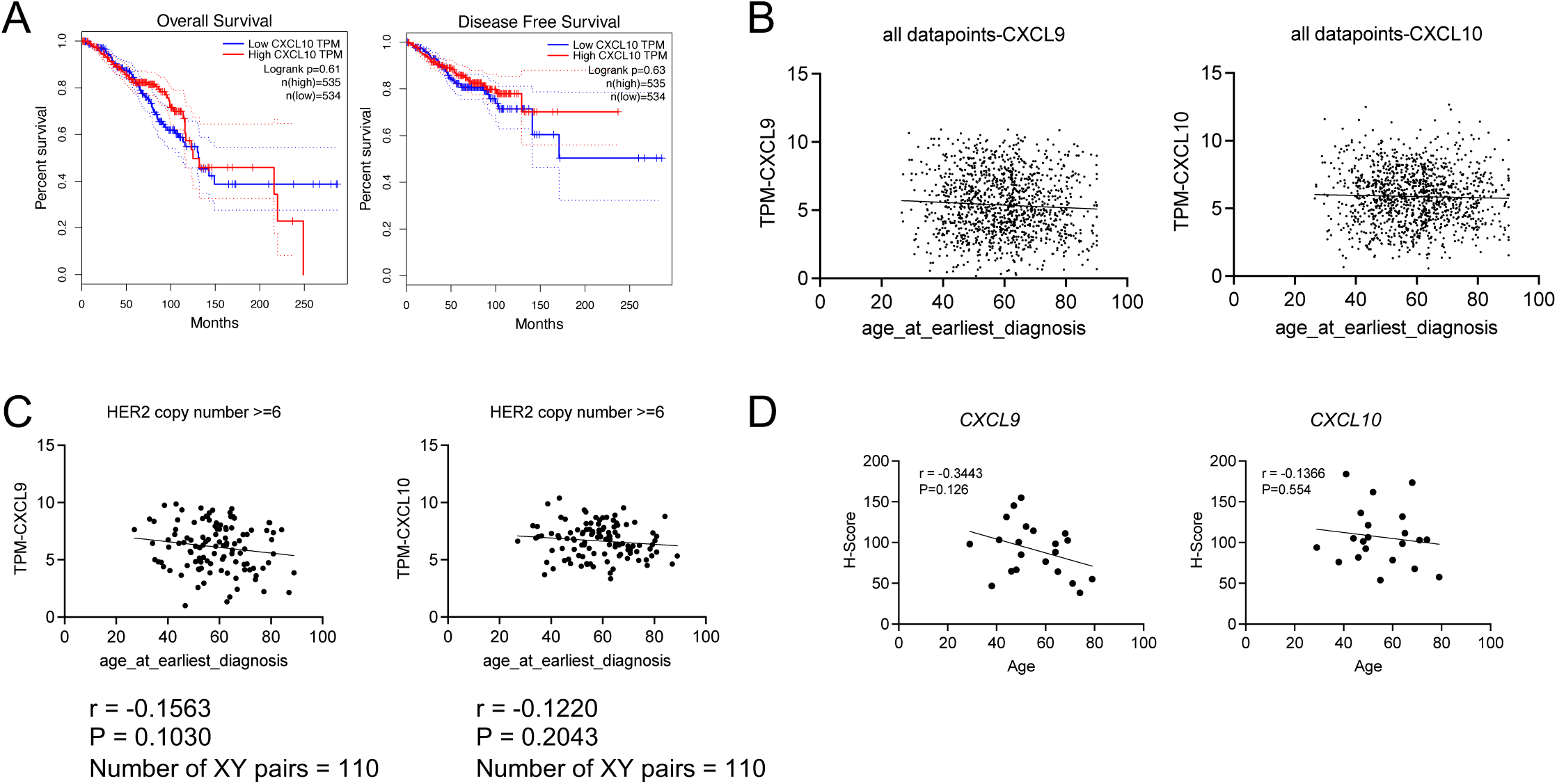
CXCL9 and CXCL10 expression with age in human breast cancer samples. Related to Figure 4. (A) Overall survival and disease-free survival curves stratified by CXCL10 expression in the GEPIA BRCA dataset. (B) The correlation between Cxcl9 and Cxcl10 RNA level with age at earliest diagnosis from TCGA BRCA dataset. (C) The correlation between *CXCL9* and *CXCL10* RNA level with age at earliest diagnosis from UCSC Xena BRCA dataset with HER2 copy number above 6. (D) *CXCL9* and *CXCL10* expression measured by RNAscope H-score from mixed breast cancer patient samples across different ages.

**Extended Data Fig. 6.**
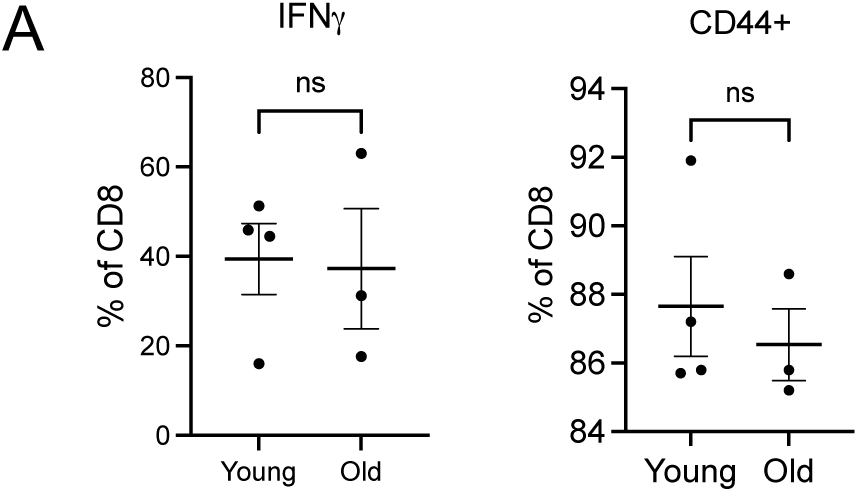
T cell in vitro activation assay. Related to Figure 6. (A) The percentage of CD8⁺ T cells expressing IFNγ or CD44. Each dot represents one mouse. Unpaired t-test was used for statistical analysis. ns P >= 0.05. * P<0.05.

**Extended Data Fig. 7.**
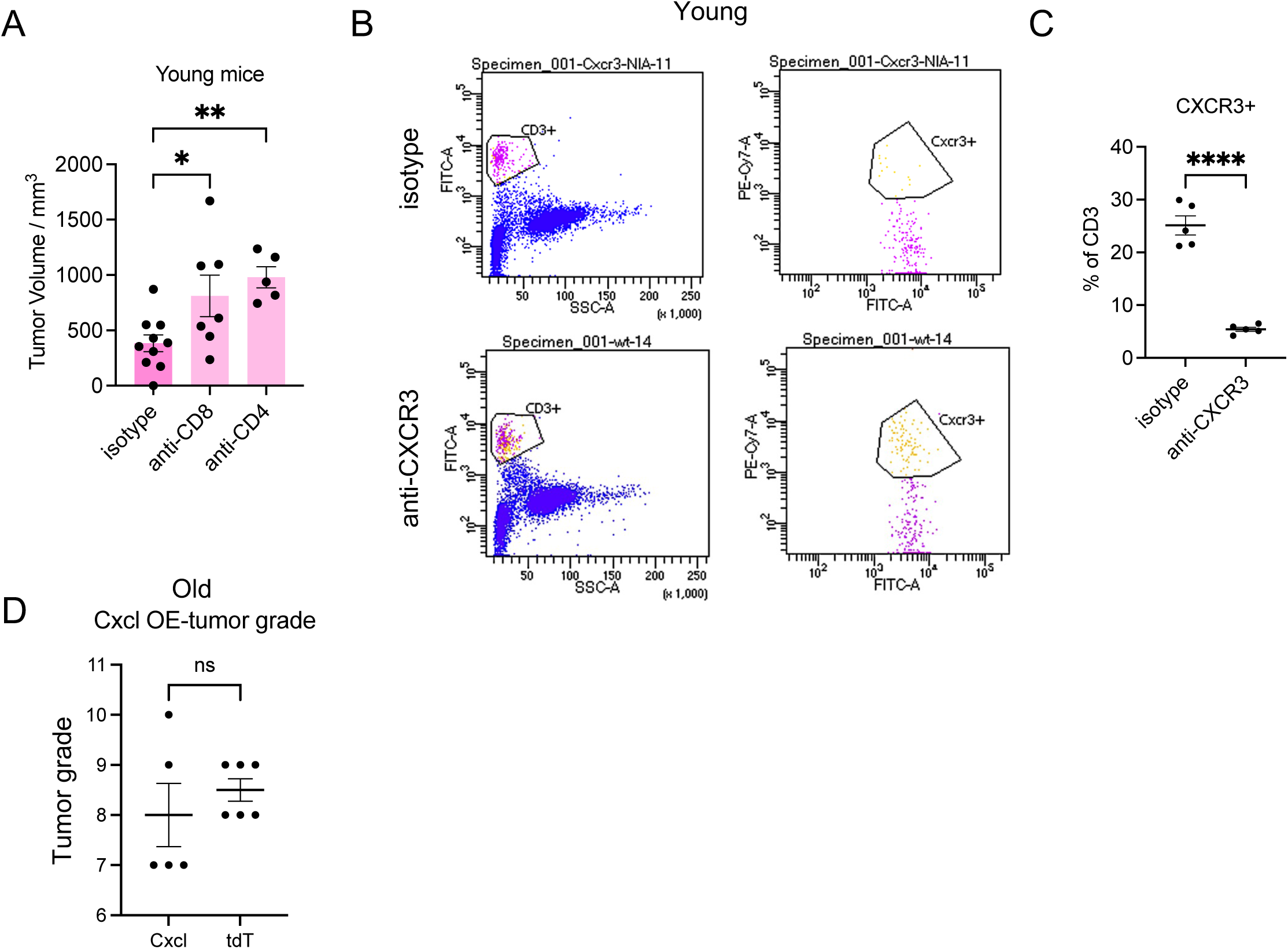
Functional test of CXCL9/10 – T cell axis in young and old mice. Related to Fig 7. (A) Tumor volumes in young mice treated with isotype or anti-CD4 or an-CD8 antibodies were measured 15 days after tumor induction. (B) Representative FACS gating results of CD3 and CXCR3 positive cells from mice treated with isotype or CXCR3 antibodies. (C) The percentage of CD3+ T cells expressing CXCR3. (D) Tumor grade assessed from mammary glands 15 days after lenti-HER2-Cxcl9-Cxcl10 or lenti-HER2-tdTomato injection. For (A), ANOVA was used for statistical analysis. For (C,D), unpaired t-test was used for statistical analysis. ns P >= 0.05. * P<0.05. ** P<0.01. **** P<0.001.

